# IL-10 receptor blockade delivered simultaneous with BCG vaccination sustains long term protection against *Mycobacterium tuberculosis* infection in mice

**DOI:** 10.1101/2021.09.05.458995

**Authors:** Varun Dwivedi, Shalini Gautam, Colwyn A. Headley, Tucker Piergallini, Jordi B. Torrelles, Joanne Turner

## Abstract

*Mycobacterium bovis* bacillus Calmette-Guérin (BCG) immunization still remains the best vaccination strategy available to control the development of active tuberculosis (TB). Protection afforded by BCG vaccination gradually wanes over time and while booster strategies have promise, they remain under development. An alternative approach is to improve BCG efficacy through host-directed therapy. Building upon prior knowledge that blockade of interleukin-10 receptor 1 (IL-10R1) during early *Mycobacterium tuberculosis* (*M.tb*) infection improves and extends control of *M.tb* infection in mice, we employed a combined anti-IL-10R1/BCG vaccine strategy. A subcutaneous, single vaccination of BCG/αIL10-R1 increased the numbers of CD4^+^ and CD8^+^ central memory T cells, and reduced TH1 and TH17 cytokine levels in the lung for up to 7 weeks post vaccination. Subsequent *M.tb* challenge in mice showed both an early (4 week) and sustained long-term (47 week) control of infection, which was associated with increased survival. In contrast, protection of BCG/saline vaccinated mice waned 8 weeks post *M.tb* infection. Our findings demonstrate that a single and simultaneous vaccination with BCG/αIL10-R1 sustains long-term protection, identifying a promising approach to enhance and extend the current BCG mediated protection against TB.

## INTRODUCTION

Tuberculosis (TB), caused by *Mycobacterium tuberculosis (M.tb*) is considered a global public health emergency, as declared by the World Health Organization (WHO) in 1993. TB is a leading cause of mortality, associated with more than 10 million new cases and 1.2 million deaths each year (1). The ongoing TB pandemic is worsened by the emergence of multi-drug resistance strains, opportunistic co-infections, and limited advancements in chemotherapeutics.

Vaccination is considered the most potent and cost-effective approach for improving public health in both the industrialized and developing world (2–4). Currently *Mycobacterium bovis* bacillus Calmette-Guérin (BCG) is the only available vaccine to provide immune protection against TB (5–8). The BCG vaccine elicits a robust TH1 response, which is critical towards mitigating TB burden(9–12). Despite this, BCG has limited protection against pulmonary TB in adults in high TB-endemic countries and is estimated to prevent only 5 to 15% of all potentially vaccine preventable deaths due to TB (1, 13, 14). Therefore, BCG-based vaccine strategies should aim at inducing long lasting T cell responses that would contribute to long-term protection.

Interleukin 10 (IL-10) is a correlate of TB disease in mice and humans (15–18). We and others have shown that IL-10 negatively regulates the immune response during *M.tb* infection in both *M.tb-*resistant C57BL/6 and *M.tb-*susceptible CBA/J mice (16, 19–21). We have also shown that *M.tb* infection of CBA/J mice leads to increased IL-10 in the lungs during the later phase of infection and that blockade of IL-10 receptor 1 (IL-10R1) at that time, promotes TH1 immunity and stabilizes the bacterial burden (21). Important to this study, IL-10R1 blockade during the first 21 days of *M.tb* infection, when IL-10 levels in the lung are negligible, resulted in early recruitment of TH1 cells to the lung and improved long-term control of *M.tb* for at least 120 days post-infection. This protection was associated with the formation of mature fibrotic granulomas and extended host survival (22).

While early IL-10R1 blockade can substantially improve long-term control of *M.tb* infection, changes in *M.tb* control were not evident until 60 days post-infection (22). We therefore hypothesize that IL-10 negatively influences the initial generation of adaptive immunity required for long lasting control of *M.tb* infection. This concept is supported by previous studies showing that IL-10 can directly or indirectly restrict memory CD4^+^ and CD8^+^ T cell differentiation (23–27). Studies have also shown that IL-10R1 blockade for 3-weeks following BCG vaccination could enhance antigen specific TH1 and TH17 immune responses in the lungs, that subsequently reduced the *M.tb* burden for up to 16 weeks post infection (28) showing proof of concept for our hypothesis. Here, we specifically asked whether a single dose of IL-10R1 antibody, delivered simultaneous with the BCG vaccine, was sufficient to enhance the long-term control of *M.tb* infection afforded by BCG. This strategy spatially separates the influence of IL-10 on the generation of protective immunity to BCG from the impact of IL-10 on control of *M.tb* infection. Our results reveal that a single dose of αIL-10R1 delivered simultaneous with BCG vaccination stimulated immunity that was capable of maintaining long-term control of *M.tb* infection. Long-term protection was sustained for at least 47 weeks post *M.tb* infection, and was associated with extended survival relative to mice receiving BCG alone. Prior to *M.tb* challenge, a single αIL-10R1/BCG vaccination resulted in the emergence of CD4^+^ and CD8^+^ central memory T cells, a reduced pro-inflammatory cytokine profile in the lungs, and increased *M.tb* antigen specific IFN-γ and IL-17 production. These findings identify IL-10 as an important immuno-modulator that impedes the development of long-term BCG specific memory immunity. Our studies also demonstrate that a single dose of αIL-10R1 delivered simultaneous with BCG vaccination is sufficient to reverse the known waning protective efficacy of BCG. Furthermore, our results identify a single-dose αIL-10R1 strategy that may be more amendable to implement in humans.

## MATERIALS AND METHODS

### Mice

Six to eight-week-old, specific pathogen free male or female wild type CBA/J strain mice (WT) were purchased from The Jackson laboratory (The Jackson Laboratory, Bar Harbor, ME). Mice were acclimatized for at least 1 week before any experimental manipulation. IL-10^−/−^ mice on the CBA/J strain background were developed and bred in house (stock now available at The Jackson Laboratories; strain 036145 CBA.129P2(B6)-Il10<tm1Cgn>/TrnrJ) and age and sex matched with WT mice. Mice were housed in ventilated micro isolator cages in ABSL-2 or ABSL-3 animal facilities and maintained with sterilized water and food *ad libitum*. Mice were euthanized at predetermined time points by CO_2_ asphyxiation. The Ohio State University (2009A0226) or Texas Biomedical Research Institute (1617 MU) Institutional Laboratory Animal Care and Use Committees approved animal protocols.

### Immunization

BCG Pasteur (ATCC 35734) and *M.tb* Erdman (ATCC 35801) were grown in supplemented Proskauer-Beck medium as previously described (29). Mice were immunized by the subcutaneous (s.c.) route with 1×10^5^ colony forming units (CFU) of BCG prepared in normal saline (0.9% NaCl) or with normal saline alone (sham). To co-immunize mice, 1×10^5^ BCG and 1.6 mg of αIL-10R1 antibody (BE0050, clone: 1B1.3A; BioXcell) or its isotype IgG1 control antibody (BE0088, clone: HRPN; BioXcell) were admixed prior to vaccination. All vaccine and control formulations were given in a 200 μl final volume. IL-10 knock-out (IL-10^−/−^) mice were immunized with 1×10^5^ CFU of BCG in normal saline (0.9% NaCl) or normal saline alone (sham) by the s.c. route.

### *M.tb* infection and determination of bacterial load

Seven weeks post-BCG immunization, mice were infected with a low dose aerosol of *M.tb* Erdman using a Glass-Col inhalation exposure system (Terre Haute, IN) calibrated to deliver 50-100 CFU to each individual mouse (29). Mice were euthanized at specific time points post-infection and lungs were aseptically harvested and homogenized. Lung homogenates were serially diluted and plated onto 7H11 agar plates enriched with OADC (oleic acid, albumin, dextrose, and catalase, Sigma-Aldrich, St. Louis, MO) and incubated at 37ºC for 3 weeks. CFU were counted to determine the burden in each organ (30). Mice allocated to survival studies were monitored over a period of 50 weeks, and any surviving mice were euthanized at study endpoint. Mice were euthanized when they reached a body condition score of 2 or less (31). Scores were determined by weekly visual and hands-on examination of each animal.

### Lung mononuclear cell isolation

Mice were euthanized at pre-determined time points post vaccination or *M.tb* challenge and lungs perfused with 10 ml PBS containing 50 U/ml heparin via the right ventricle of the heart. Lung lobes were extracted and placed in enriched complete Dulbecco’s modified Eagle’s medium [DMEM containing L-glutamine (Life Sciences, Tewksbury, MA, USA)], supplemented with sterile-filtered mixture of 5 ml HEPES buffer (1 M; Sigma), 10 ml MEM nonessential amino acid solution (100X; Sigma), 660 μl 2-mercaptoethanol (50 mM; Sigma), and 45 ml heat-inactivated FBS (Atlas Biologicals, Ft. Collins, CO, USA). Lungs were dissociated using a GentleMACS Dissociator, in the presence of collagenase A (type XI; 0.7 mg/ml obtained from *Clostridium hystolyticum*; Sigma) and type IV bovine pancreatic DNAse (30 μg/ml; Sigma) and incubated for 30 min at 37ºC, 5% CO_2_. The enzymatic reaction was stopped by adding complete DMEM. Single cell suspensions were achieved by passing the digested lung tissue through 70-μm cell strainers. Cells were treated with Gey’s solution (8 mM NH_4_Cl, 5 mM KHCO_3_ in water) to lyse residual red blood cells and suspended in complete DMEM. Live cells were counted by using a Trypan blue live-dead exclusion method or using a Cellometer K2 (Nexcelom Bioscience, Lawrence, MA) with acridine orange (AO)/propidium iodide (PI) stain (29).

### *M.tb* antigen specific cell culture

Lung mononuclear cells were cultured with medium or 10 μg/ml of *M.tb* culture filtrate proteins (CFP) for 48 h at 37°C, 5% CO_2_ (29). Culture supernatants were collected and stored at −80°C.

### Enzyme linked immunosorbent assay (ELISA)

Clarified lung homogenates and cell culture supernatants were thawed and analyzed for IFN-γ, IL-12p70, TNF-α, IL-17 and IL-10 by ELISA following the manufacturer instructions (BD Biosciences, San Jose, CA).

### Flow cytometry

Lung mononuclear cells were suspended in incomplete RPMI medium (Sigma-Aldrich) containing 0.1% sodium azide. Surface marker staining was performed as described (22). Specific antibodies for surface marker staining were purchased from Biolegend [PerCP anti-CD4 (clone: GK1.5), APC/Cyanine7 anti-CD8 (clone: 53-6.7), PE/Cy7 anti-CD62L (clone: MEL-14), APC anti-CD44 (clone: IM7), Brilliant Violet 421 anti-CD197 (CCR7; clone: 4B12)]. Briefly, cells were blocked with mouse Fc block (clone: 2.4G2; BD Biosciences) for 10 min followed by staining with fluorescent dye conjugated antibodies specific to surface markers for 20 min at 4°C in the dark. Cells were fixed and samples were acquired using a BD Canto or Beckman Coulter CyAn ADP flow cytometer and results analyzed using FlowJo software vr. 10.5 &10.6 (Tree Star, Ashland, OR).

### Histology

The right caudal lung lobe was isolated from individual mice, inflated with and stored in an excess of 10% neutral-buffered formalin as described (32). Lung tissue was processed, sectioned, and stained with hematoxylin and eosin (H&E) for light microscopy with lobe orientation designed to allow for maximum surface area of each lobe to be seen. Sections were examined in a blinded manner by a board-certified veterinary pathologist without prior knowledge of the experimental groups and evaluated according to the percent affected tissue area, granuloma distribution, granuloma character, granuloma border and cellular composition. Percent affected area was microscopically quantified by calculating the total area of the involved tissue over the total area of the lobe for each individual mouse and graded as 1, 2, 3, 4 and 5, corresponding to <10%, 10% to 25%, 25% to 50%, 50% to 75%, >75% affected tissue, respectively.

### Statistical analysis

Statistics were performed using Prism vr. 7 Software (GraphPad Software, San Diego, CA). Unpaired, two-tailed Student’s *t*-test was used for two group comparisons. Log-rank test was used to determine statistical significance of survival experiments. Statistical significance was reported as *P<0.05, **P<0.01, ***P<0.005 or ****P<0.001.

## RESULTS

### BCG vaccination stimulates IL-10 production in the lung, which is blocked by αIL-10R1

WT mice were immunized with BCG (or saline control) by the s.c. route and IL-10 levels were measured in the lung at weeks 1, 2, 4, 5, and 7 post-immunization. BCG immunization resulted in significantly higher IL-10 levels at each time point tested. IL-10 could be detected in the lung as early as 1-week post BCG vaccination and remained high for up to 7 weeks post immunization compared to the saline vaccinated group (**Fig. 1A**). These data demonstrate that s.c. BCG vaccination can stimulate IL-10 production in the lung.

**Figure 1.**
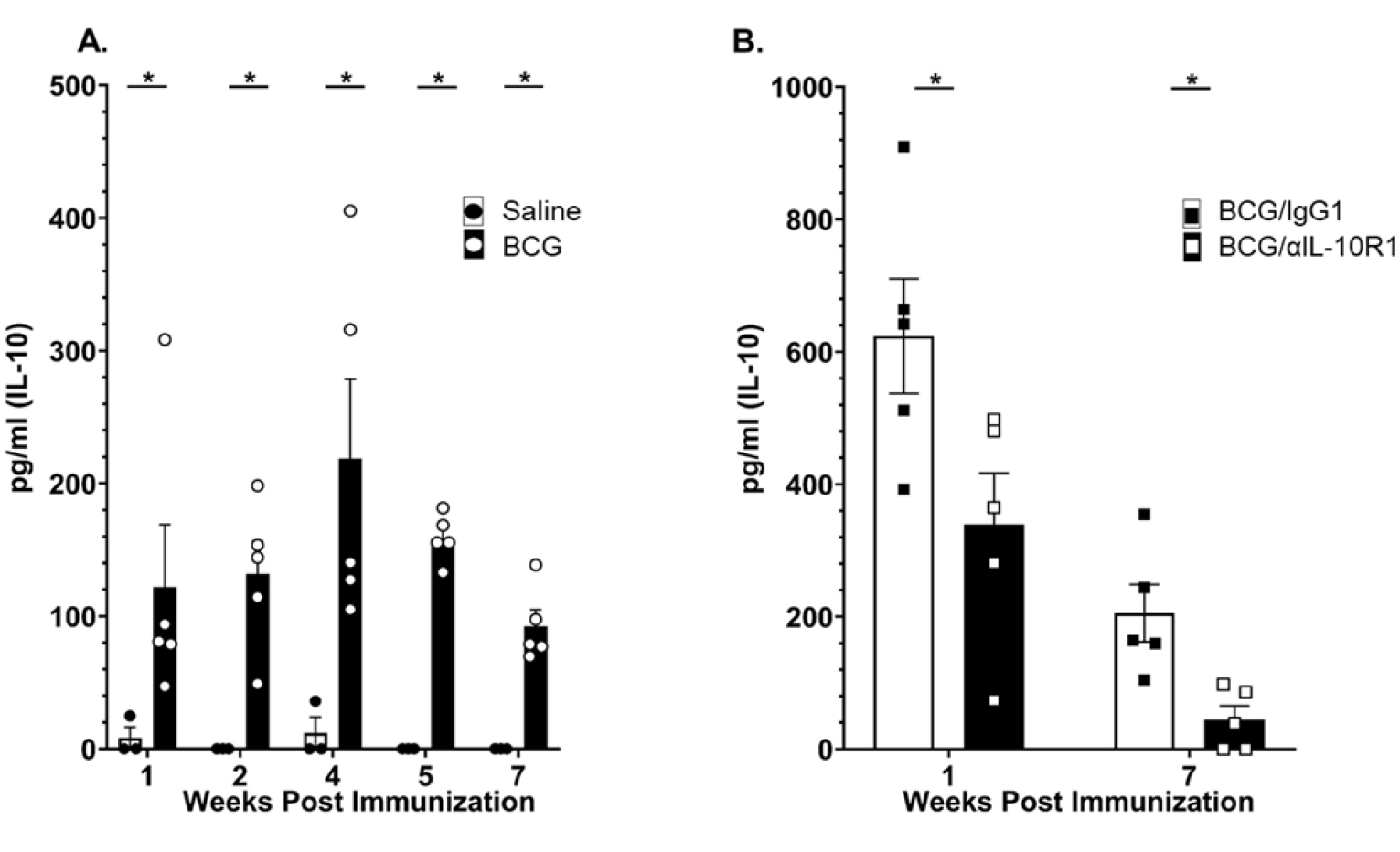
IL-10 in the lung in response to BCG or BCG/αIL10-R1 administration. WT mice were immunized with saline or BCG by the subcutaneous route and euthanized at pre-determined time points (**A**), or immunized with BCG/αIL-10R1 or BCG/IgG1 and euthanized at 1- or 7-weeks post immunization (**B**). IL-10 was determined in lung homogenate by ELISA. Data represent the mean ± SE of one of two independent experiments with 3-5 mice in each group at each time point. Student’s *t* test was performed to determine statistical significance between the saline and BCG immunized groups (A) or between the BCG/IgG1 and BCG/αIL-10R1 groups (B). *P<0.05; **P<0.001.

We determined the impact of simultaneous immunization of BCG with αIL-10R1 on the production of IL-10 in the lung. Mice were immunized with a combined dose of BCG/αIL-10R1 or BCG/IgG1 control antibody via the s.c. route. BCG/α-IL-10R1 significantly reduced IL-10 in the lung at week 1 and week 7 post immunization (Fig. 1B). These results indicate that a transient blockade of IL-10R1 delivered simultaneous with BCG vaccination has a long-term impact on local IL-10 production in the lung. IL-10 has a known deleterious impact on the control of *M.tb* infection (15) and therefore the IL-10 inducing properties of BCG vaccination, or any other lung insult that drives IL-10 production, could inadvertently have a negative consequence on *M.tb* infection control.

BCG/αIL-10R1 vaccination generated a relatively low pro-inflammatory environment in the lung compared to BCG/IgG1 vaccination. This is contrary to expectations from prior studies with *M.tb* (16, 22, 28). Indeed, we detected a significant reduction in IFN-γ, TNFα, IL-17 and IL-12p70 in the BCG/αIL-10R1 group compared to BCG/IgG1 at both time points studied (**Fig. 2A-D**). Cytokine levels in mice receiving BCG/αIL-10R1 were marginally elevated relative to mice receiving no BCG (data not shown).

**Figure 2.**
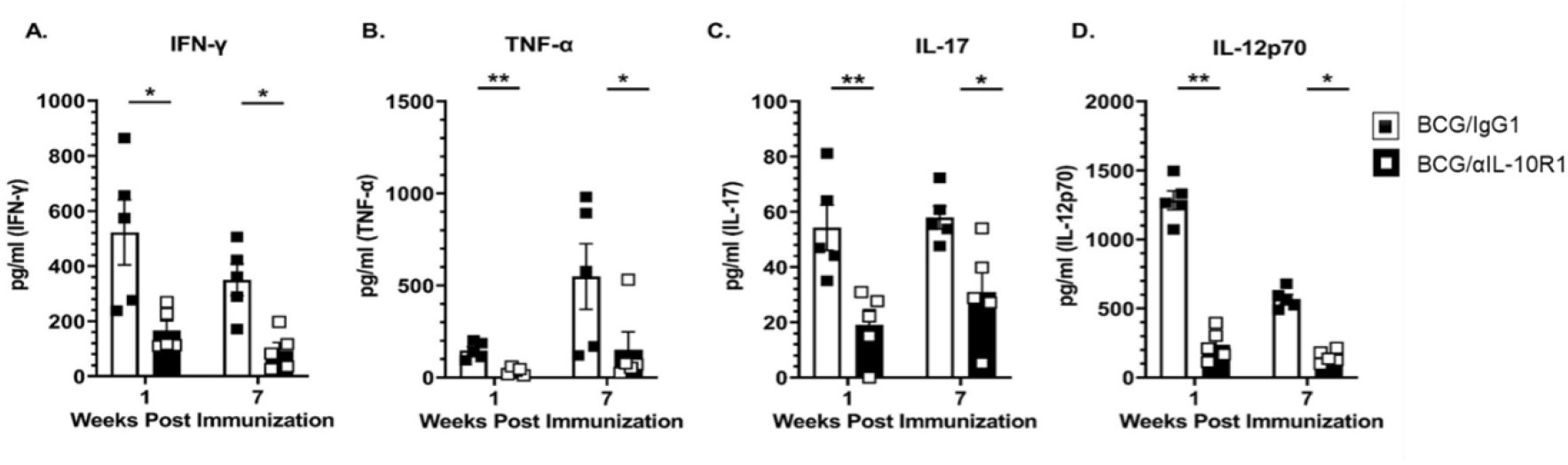
Pro-inflammatory cytokines in the lung in response to BCG/αIL-10R1 administration. WT mice were immunized with BCG/αIL-10R1 or BCG/IgG1 antibody subcutaneously. Immunized mice were euthanized at 1- and 7-weeks post immunization. Lungs were homogenized and centrifuged to obtain clarified homogenate. IFN-γ (**A**), TNF-α (**B**) IL-17 and IL-12p70 (**D**) were measured by ELISA. Data represent the mean ± SE of one of two independent experiments with 3-5 mice in each group at each time point. Student’s *t* test was performed to determine statistical significance between the BCG/IgG1 and BCG/αIL-10R1 experimental groups at 1- and 7-weeks post vaccination. *P<0.05; **P<0.01.

### BCG vaccination delivered simultaneous with αIL-10R1 generates central memory T cells in the lung

IL-10 is known to interfere with the generation of memory T cells (23–26). We therefore determined whether BCG/αIL-10R1 vaccination produced a phenotypically different memory T cell pool in the lung. The total number of CD4^+^ and CD8^+^ T cells were modestly increased 7 weeks post-immunization in mice receiving BCG/αIL-10R1 although this did not reach significance (**Figs. 3A, E**). BCG/αIL10R1 vaccination led to a consistent increase in the total number of memory (CD44^hi^), central memory (CD4^+^CD44^hi^CD62L^+^CCR7^+^) and effector memory (CD4^+^CD44^hi^CD62L^−^CCR7^−^) cells in both CD4^+^ and CD8^+^ T cell subsets, but again this did not reach significance (**Figs. 3B-D & F-H**). These results suggest that short-term IL-10R1 blockade during BCG vaccination results in a modest accumulation of central and effector memory cells at 7 weeks post vaccination in the lung.

**Figure 3.**
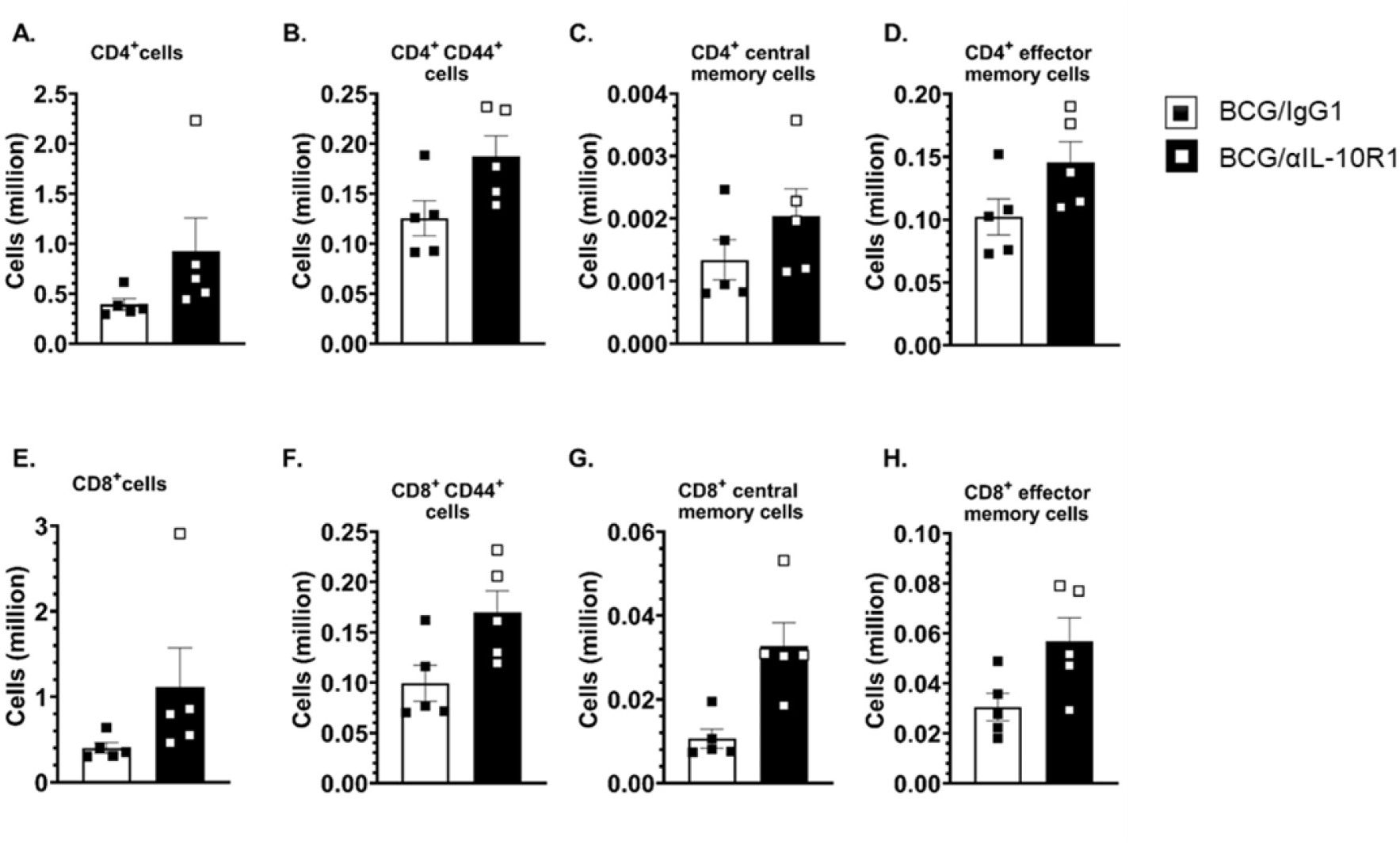
BCG/αIL-10R1 administration causes accumulation of central memory cells in lung. WT mice were subcutaneously immunized with BCG/αIL-10R1 or BCG/IgG1. Immunized mice were euthanized at 7 weeks post immunization and lung mononuclear cells were harvested and stained with fluorescent dye tagged antibodies specific for CD4 and CD8 in combination with CD44, CD62L and CCR7 and acquired by flow cytometry and analyzed by FlowJo software. Absolute number of (**A**) CD4^+^, (**B**) CD4^+^ CD44^hi^ (**C**) CD4^+^ CD44^hi^ CD62L^+^ CCR7^+^ central memory, CD4^+^ CD44^hi^ CD62L^−^ CCR7^−^ effector memory, (**E**) CD8^+^, (**F**) CD8^+^ CD44^hi^, (**G**) CD8^+^ CD44^hi^ CD62L^+^ CCR7^+^ central memory, (**H**) CD8^+^ CD44^hi^ CD62L^−^ CCR7^−^ effector memory. Data represent the mean ± SE of one of two independent experiment with 3 to 5 mice in each group. Student’s *t* test was performed to determine the statistical significance between BCG/IgG1 and BCG/α-IL-10R1 experimental groups. No statistical significant differences were found.

### BCG vaccination delivered simultaneous with αIL-10R1 increases antigen specific Th1 and Th17 cytokine production by lung cells

IL-10 suppresses TH1 and TH17 immune responses necessary to control *M.tb* infection (28, 33–37). Additionally, BCG/αIL-10R1 vaccination reduced IL-10 levels (**Fig. 1B**) and stimulated a moderate accumulation of memory T cells in the lung (**Fig. 3C, G**). We therefore determined whether BCG/αIL-10R1 vaccination could enhance antigen specific TH1 and TH17 cytokine production. Lung cells from WT mice receiving BCG/αIL-10R1 secreted significantly more antigen specific IFN-γ, TNF-α, and IL-17 (**Fig. 4A-C**) than those receiving BCG/IgG1. Therefore, IL-10 receptor blockade at the time of BCG vaccination promoted the generation of functional TH1 and TH17 antigen specific T cells in the lungs.

**Figure 4.**
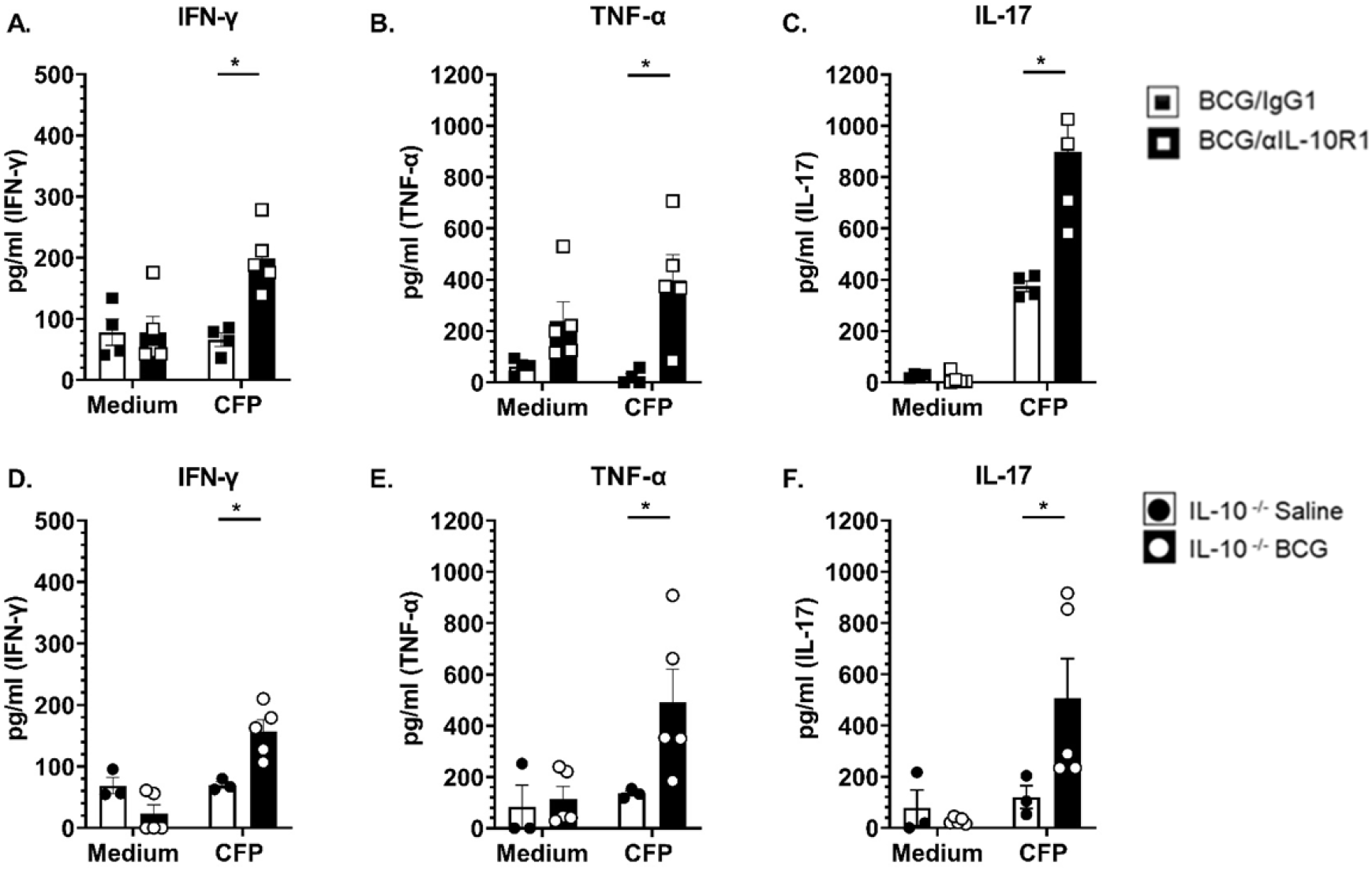
BCG/αIL-10R1 administration increases antigen specific Th1 and Th17 cytokine responses. WT mice were immunized with BCG/αIL-10R1 or BCG/IgG1 (**A-C**). IL-10 ^−/−^ CBA/J mice were vaccinated with saline or BCG (**D-F**). Mice were euthanized at 7 weeks post vaccination and lung mononuclear cells *ex-vivo* stimulated without (medium) or with CFP for 48 hrs. Culture supernatant were analyzed for the production of IFN-γ (**A, D**), TNF-α (**B, E**) and IL-17 (**C, F**) respectively. Student’s *t* test was performed to determine statistical significance between the response to CFP of BCG/IgG1 and BCG/αIL-10R1 experimental groups (A-C) or IL-10 ^−/−^ saline and BCG experimental groups (D-F). *P<0.05.

To further establish that the absence of IL-10 signaling during BCG vaccination increases antigen specific TH1 and TH17 cytokine secretion, we immunized IL-10^−/−^ mice with BCG or saline (control) and assessed antigen specific cytokine responses by lung cells 7 weeks post vaccination. Similar to WT mice vaccinated with BCG/αIL10R1, IL-10^−/−^ mice vaccinated with BCG alone had significantly increased production of antigen specific IFN-γ, TNFα, and IL-17 (**Fig. 4D-F**), compared to IL-10^−/−^ mice immunized with saline. These results suggest that the absence of IL-10, similar to our data blocking the action of IL-10R1, promotes a population of T cells capable of secreting TH1 and TH17 cytokines in the lungs.

### BCG vaccination delivered simultaneous with αIL-10R1 provides long term protection against *M.tb* infection

WT mice were vaccinated subcutaneously with BCG/αIL-10R1, BCG/IgG1 or saline only. In parallel, IL-10^−/−^ mice were vaccinated with BCG or saline. Seven weeks post-immunization, mice were challenged with a low dose aerosol *M.tb* infection. As previously reported, IL-10^−/−^ mice were capable of reducing *M.tb* bacterial burden in the lung to a greater extent than WT mice (22) (**Fig. 5A** saline *vs*. **Fig. 5B** saline). BCG vaccination further reduced the bacterial burden in IL-10^−/−^ mice by 1-Log_10_ at 4 weeks post-infection, and importantly, this reduction was sustained for at least 52 weeks post infection (**Fig. 5A**).

**Figure 5.**
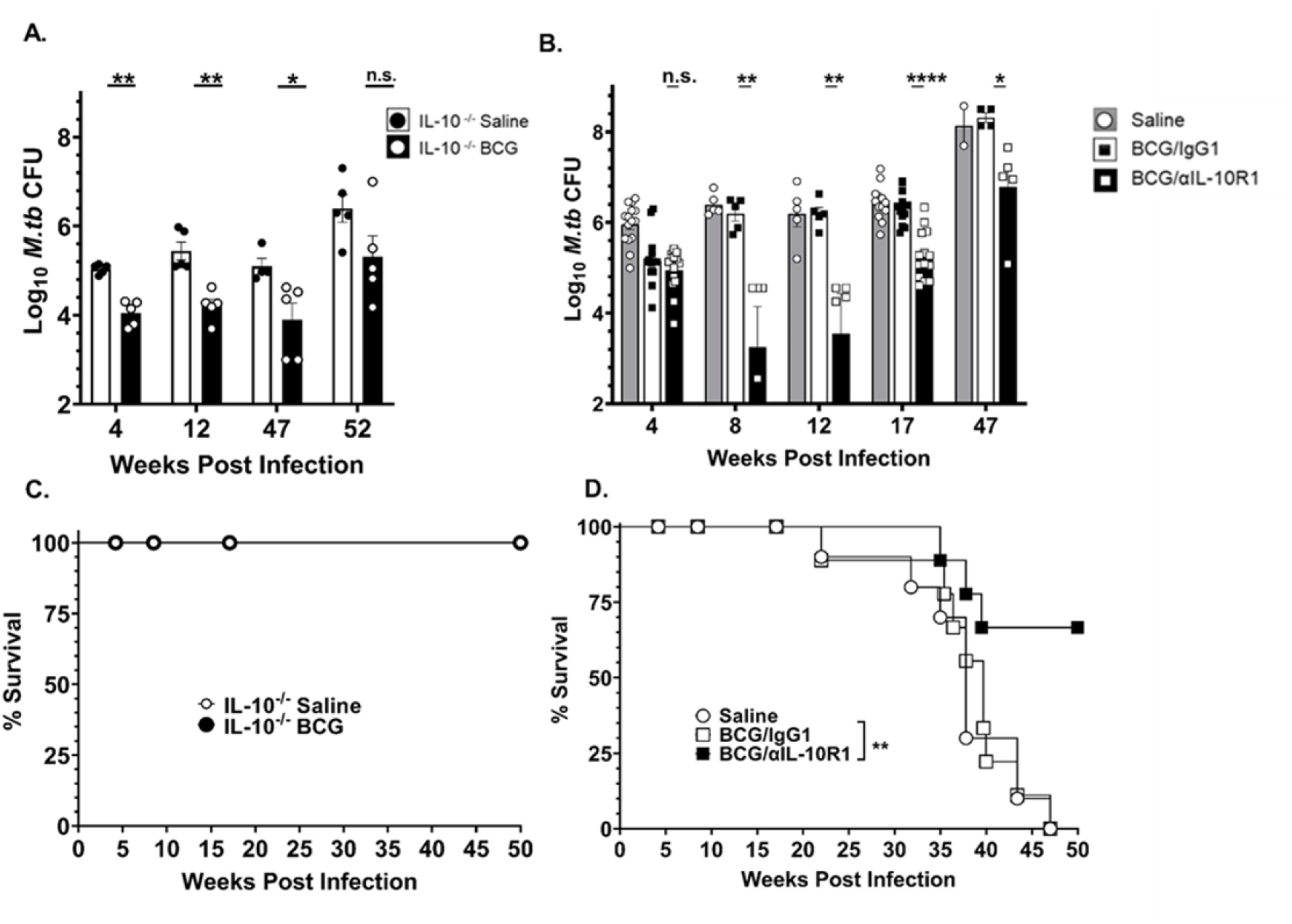
BCG/αIL-10R1 administration provides long-term protection against *M.tb* infection. IL-10^−/−^ CBA/J mice were vaccinated with saline or BCG (**A, C**). WT mice were immunized with saline or BCG/αIL-10R1 or BCG/IgG1 (B, D). Mice were infected with *M.tb* at 7-week post vaccination. Lung CFU counts in IL-10^−^/^−^ (**A**) or WT (**B**). Survival curve for IL-10^−/−^ (**C**) and WT (**D**) mice. Data in figure 5B are a combined 1 to 3 independent experiments each having 5 mice in each group in all data points. Student’s *t* test was performed to determine statistical significance between BCG/α-IL-10R1 and BCG/IgG1 (WT) or between saline and BCG (IL-10^−/−^) immunized experimental groups. *P <0.05; **P<0.01; and P ****P<0.0001. Data in C and D represent a single experiment with 10-14 mice in each group. Log-rank test was used to determine statistical significance of survival between BCG/αIL-10R1 and BCG/IgG1.

In WT mice, the BCG/αIL-10R1 and BCG/IgG1 vaccinated groups both significantly reduced the *M.tb* bacterial burden by approximately 1-Log_10_ at week 4 post *M.tb* infection compared to the unvaccinated (saline) group (**Fig. 5B**). As expected, WT mice that received BCG/IgG1 vaccination quickly lost protection afforded by BCG vaccination alone (37, 38), with lung bacterial burden resembling unvaccinated mice as early as 8-week and reaching almost 8 Log_10_ CFUs in the lung by 47-week post infection (**Fig. 5B**). In contrast, WT mice vaccinated with BCG/αIL-10R1 further reduced the *M.tb* burden at week 8 post infection, which was sustained at a significantly lower CFU for up to 47 weeks post infection, albeit at slowing increasing levels (study end point, **Fig. 5B**).

An independent survival study was established for BCG or saline vaccinated IL-10^−/−^ mice (**Fig. 5C**) and for BCG/αIL-10R1, BCG/IgG1 or saline vaccinated WT mice (**Fig. 5D**). BCG or saline vaccinated IL-10^−/−^ mice (**Fig. 5C**) challenged with *M.tb* showed that the complete absence of IL-10 resulted in 100% survival at least to 50 weeks in both BCG or saline vaccinated mice, demonstrating the detrimental impact of IL-10 throughout *M.tb* infection as we described (16, 21).

While we observed no survival advantage for BCG vaccination in IL-10^−/−^ mice, extrapolation of bacterial burden (**Fig. 5A**) and pathology scores (**Fig. 6D**) suggests that BCG vaccination would likely have extended survival further relative to IL10^−/−^ mice receiving saline. BCG/IgG1 in WT mice failed to extend survival, indicating that despite an early reduction in *M.tb* bacterial burden at week 4, BCG vaccination alone provided no survival advantage. In contrast, administration of BCG/αIL-10R1 vaccination in WT mice significantly increased survival, with over 65% of mice surviving to the study end point of 50 weeks.

**Figure 6.**
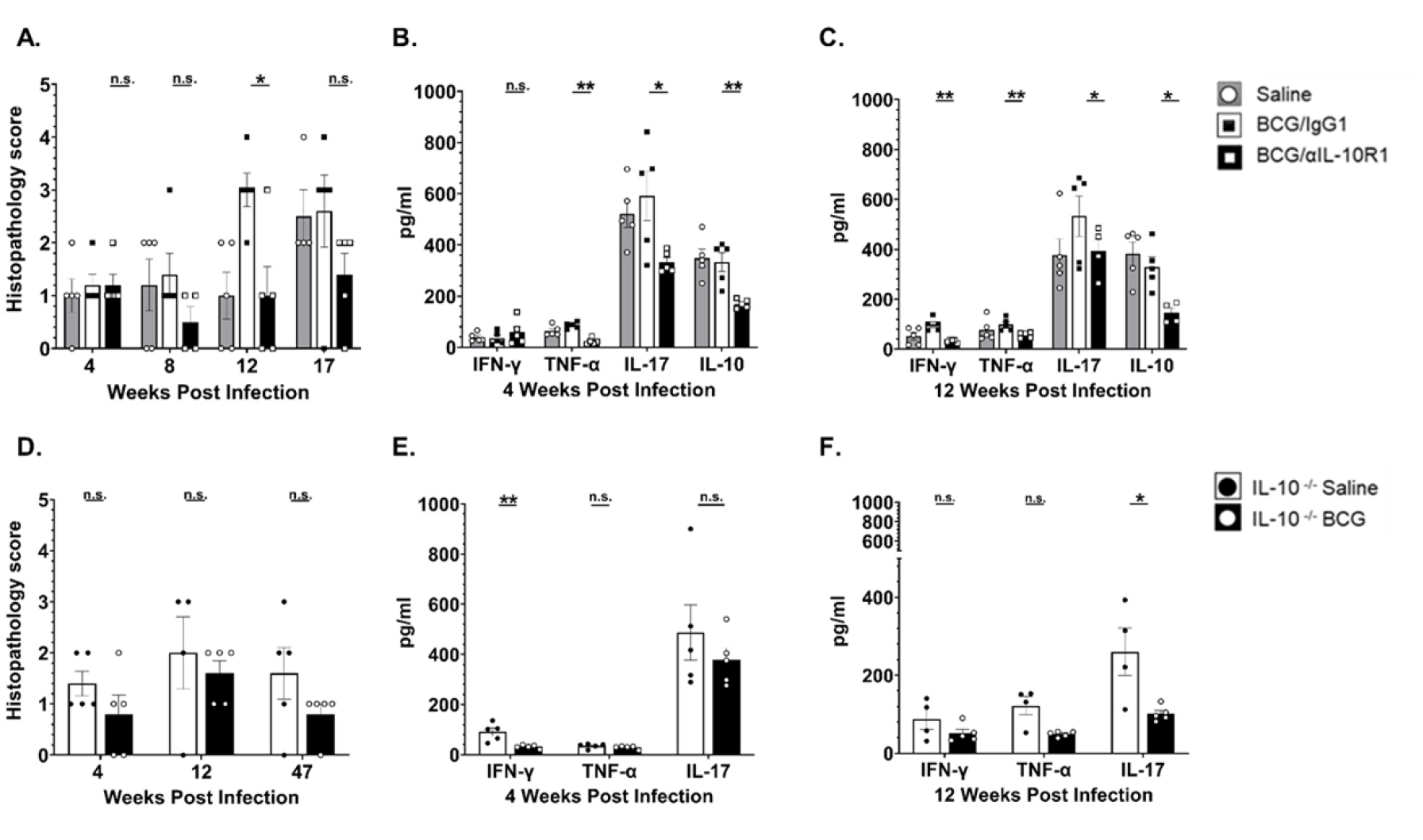
BCG/αIL-10R1 administration reduces immuno-pathology in lungs after *M.tb* infection. WT mice were immunized with saline, BCG/αIL-10R1 or BCG/IgG1 (**A-C**). IL-10^−/−^ CBA/J mice were vaccinated with saline or BCG (**D-F**). Mice were infected with *M.tb* 7-weeks post vaccination. WT and IL-10^−/−^ mice were euthanized at predetermined time points post infection and the caudal lung lobe was quantified for pulmonary inflammation as percent of tissue involved. Pulmonary inflammation in (**A**) WT mice at 4, 8, 12 and 17 weeks post infection, (**D**) IL-10^−/−^ mice at 4, 12 and 47 weeks post infection. Percent affected area was quantified by calculating the total area of the involved tissue over the total area of the lobe for each individual mouse and graded as 1, 2, 3, 4 and 5 which corresponded to <10%, <25%, 50%, <75%, >75% of affected tissue, respectively. ELISA was performed on lung homogenates to measure the level of TNF-α, IL-17, IFN-γ and IL-10 in (**B**) WT mice at 4 weeks post infection, (**C**) WT mice at 12 weeks post infection, (**E**) IL-10 ^−/−^ mice at 4 weeks post infection and (**F**) IL-10 ^−/−^ mice at 12 weeks post infection. Data in Figure 6A-C represent the mean ± SE of one of two independent experiments with 3 to 5 mice in each group at each time point. Student’s *t* test was performed to determine statistical significance between BCG/αIL-10R1 and BCG/IgG1 (WT mice) or between saline and BCG (IL-10^−/−^ mice), immunized experimental groups. *P<0.05; **P<0.01.

### BCG vaccination delivered simultaneous with αIL-10R1 reduces immuno-pathology in lungs

we also evaluated the effect of vaccination on lung pathology by assessing the degree of tissue involvement through quantification of cellular aggregation relative to the total size of the lung. All WT mice, independent of experimental groups, had similar and minimal tissue involvement (>10%) in the lung at 4 weeks post infection (**Fig. 6A**) although some pro-inflammatory cytokines in homogenates were significantly reduced in the BCG/αIL-10R1 group compared to BCG/IgG1 immunized mice at that time (**Fig. 6B**). These results indicate that even though there was no difference in *M.tb* burden or pathology, an early increased pro-inflammatory cytokine response in the BCG/IgG1 vaccinated group may augment lung pathology at later time points. This was confirmed at week 12 post infection where BCG/IgG1 vaccinated mice had abundant cellular infiltration and inflammation, with about 50% of the lung involved (**Fig. 6A**), although TH1 cytokine levels remained similar to the 4 weeks post infection time-point (**Fig. 6C**). Increased pathology scores were associated with high *M.tb* burden in the lungs of BCG/IgG1 treated mice at later time-points (**Fig. 5B**). These findings contrast with BCG/αIL-10R1 vaccinated mice that maintained a reduced lung involvement (**Fig. 6A**), correlating with less *M.tb* burden (**Fig. 5B**), which was maintained even at later time points. BCG/αIL-10R1 vaccinated mice also had significantly reduced pro-inflammatory cytokines levels (TNF-α, IL-17), as well as immunomodulatory cytokines (IL-10) in their lungs compared to the BCG/IgG1 vaccinated group at both 4- and 12-weeks post infection (**Fig. 6 B,C**). BCG/αIL-10R1 vaccinated mice also had significantly less IFN-γ in their lungs compared to the BCG/IgG1 vaccinated group at the 12-week post infection. (**Fig. 6C**).

Similar to WT mice, both saline and BCG immunized IL-10^−/−^ mice had minimal tissue involvement in the lungs at week 4 post-infection; however, BCG immunization showed reduced inflammation scores (<25%) at week 4 post-infection, which was sustained through weeks 47 post-infection (**Fig. 6D**), consistent with lower *M.tb* burden in the lung (Fig. 5A). IL-10^−/−^ mice receiving BCG showed a reduction in some TH1/pro-inflammatory cytokines in lung homogenates with a significant reduction in IFN-γ at week 4 and IL-17 at week 12 post infection as compared to saline group (**Figure 6 E-F**).

### Central memory subset of CD4^+^ and CD8^+^ T cells are associated with long term protection against *M.tb* challenge in mice vaccinated with BCG delivered simultaneous with αIL-10R1

Following vaccination with BCG/αIL-10R1 and prior to *M.tb* challenge, we observed a modest increase in CD4^+^ and in CD8^+^ central and effector memory T cells in the lung (**Fig. 3**). We therefore evaluated the same T cell subsets in the lung at week 4 and 8 post *M.tb* infection (**Fig. 7**). Results showed modest differences in total CD4^+^ and CD8^+^ T cell subsets in the lung between BCG/αIL-10R1 and BCG/IgG1 vaccinated WT mice at week 4 post *M.tb* infection (**Fig. 7A, E**). At 4 weeks post-infection, we detected a significant increase in the total number of both CD4^+^ and in CD8^+^ central memory T cells in mice that received BCG/αIL-10R1 vaccination (**Fig. 7C, G**), which was sustained through week 8 post-infection (Fig**. 7K, O**). These data suggest that a single and simultaneous vaccination with BCG/αIL-10R1 pre-*M.tb* challenge can induce central memory cells that are associated with long-term protection against *M.tb* infection. CD4^+^ and CD8^+^ effector memory T cells were detected at similar or modestly increased levels by both BCG/αIL-10R1 and BCG/IgG1 vaccinated WT mice at 4 and 8 weeks (**Fig. 7 D, H**).

**Figure 7.**
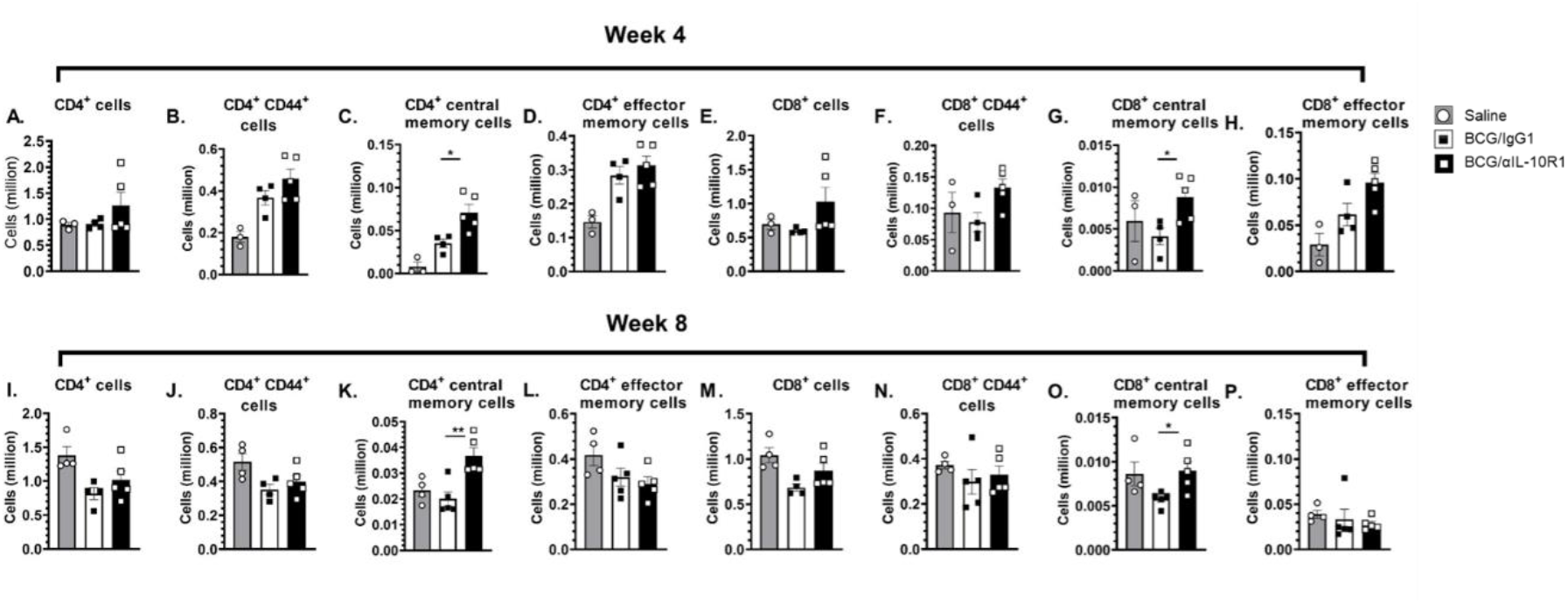
BCG/ αIL-10R1 administration enhances central memory T-cell responses in lung of *M.tb* infected mice. WT mice were subcutaneously immunized with saline, BCG/αIL-10R1 or BCG/IgG1. 7 weeks later, mice were challenged with *M.tb* and euthanized at the 4-(**A-H**) and 8-week (**I-P**) post infection. Figure represents absolute number of (A, I) CD4^+,^ (B,J) CD4^+^ CD44^hi^, (C,K) CD4^+^CD44^hi^CD62L^+^CCR7^+^ central memory, (D,L) CD4^+^CD44^hi^CD62L^−^CCR7^−^ effector memory, (E,M) CD8^+^, (F,N) CD8^+^CD44^hi^, (G,O) CD8^+^CD44^hi^CD62L^+^CCR7^+^ central memory, (H,P) CD8^+^CD44^hi^CD62L^−^CCR7^−^ effector memory. Data are the mean ± SE of one of the two independent experiment with 3 to 5 mice in each group. Student’s *t* test was performed to determine the statistical significance between BCG/αIL-10R1 or BCG/IgG1 experimental groups. *P<0.05; **P<0.01.

IL-17 and IFN-γ are critical for mediating immunity against *M.tb* infection as well as vaccine induced protection against the development of TB (39, 40). Lung cells from vaccinated and *M.tb* challenged mice were cultured *ex-vivo* with CFP and IFN-γ and IL-17 production was measured. Relative to unstimulated cells, antigen specific IFN-γ and IL-17 secretion was increased for both BCG/αIL-10R1, BCG/IgG1 vaccinated groups at week 4 and 8 post *M.tb* infection (**Fig. 8 A-D**). There was no significant differences between the BCG/αIL-10R1, BCG/IgG1 groups (**Fig. 8A-D**), with the exception of IL-17 at week 4 (**Fig. 8B**) although significance was lost by week 8 post *M.tb* infection (**Fig. 8D**). IL-17 secretion in unstimulated cultures remained high, perhaps representing a non-T cell source of IL-17 (**Fig. 8 B, D**).

**Figure 8.**
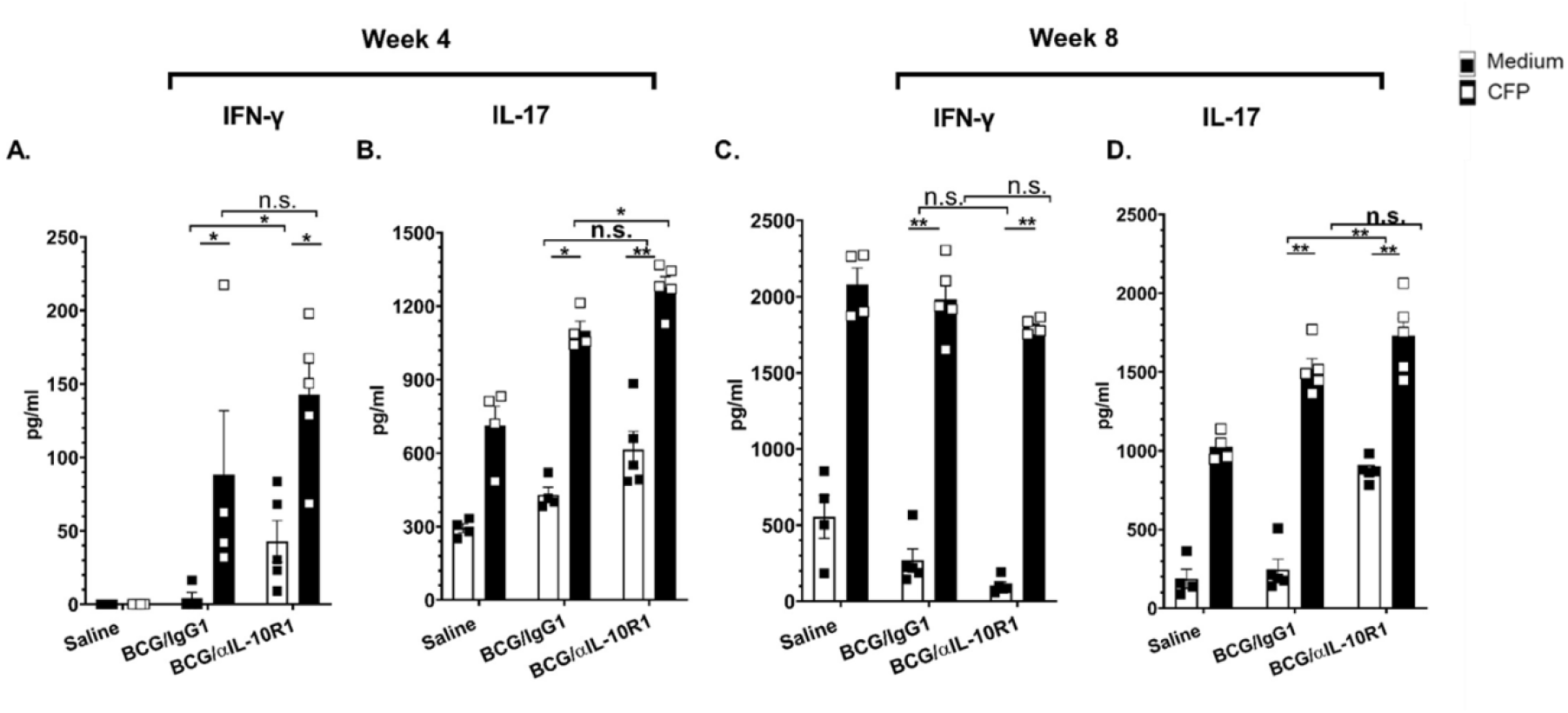
BCG/αIL-10R1 administration enhances antigen specific IFN-γ and IL-17 levels in lungs after *M.tb* infection. WT mice were s.c. immunized with saline, BCG/αIL-10R1 or BCG/IgG1. At 7 weeks later, mice were challenged with *M.tb* and euthanized at 4 and 8 weeks post infection. Isolated lung mononuclear cells were *ex-vivo* stimulated with medium or *M.tb* CFP for 48 hrs. Culture supernatant of WT mice were analyzed for the production of (**A, C**) IFN-γ and (**B, D**) IL-17 by ELISA. Student’s *t* test was performed to determine the statistical significance between the medium and CFP stimulation within an experimental group and between the groups. *P<0.05; **P<0.01.

## Discussion

In this study, we demonstrate that a single co-administration of αIL-10R1/BCG substantially improved the long term protective efficacy of BCG against *M.tb* infection for up to 47 weeks in the CBA/J mouse model, a mouse strain that is defined as relatively susceptible to *M.tb* (41, 42). We therefore demonstrate that a single dose host-directed therapy combined with BCG can improve vaccine efficacy. We also identify the specific time of BCG vaccination for a single-dose IL-10 blockade intervention to positively influence the development of long-term protective immunity against *M.tb* infection.

Our vaccination strategy was based upon prior observations from our group that early αIL-10R1 treatment of *M.tb* infected mice could substantially improve control of *M.tb* infection in CBA/J mice (22). We hypothesized that αIL-10R1 treatment blocks the negative influence of IL-10 on priming and development of long-term memory immunity. Because the presence of IL-10 can also influence maintenance of *M.tb* infection as we have described (21), we separated immune cell priming from *M.tb* infection by blocking the action of IL-10 during αIL-10R1/BCG vaccination instead of at *M.tb* infection. The superior control of *M.tb* infection that we observed in αIL-10R1/BCG vaccinated mice demonstrates that IL-10 inhibits the early development of long-term memory immunity. Further supporting our findings, we have previously demonstrated that selective delipidation of BCG, which induces less IL-10, generates enhanced protection against *M.tb* challenge minimizing tissue damage when compared to conventional BCG (43).

IL-10R1 blockade during vaccination has been tested in various short term experimental or disease models (44–46), including a mouse model of herpes virus (HPV) 16 E7 transformed TC-1 tumor growth, where inclusion of αIL-10R with vaccination enhanced specific cytolytic (CTL) responses (47). Similarly, blockade of IL-10R1 permitted an otherwise ineffective DNA vaccine to become highly efficient at stimulating CD4^+^ and CD8^+^ T cell responses, leading to accelerated clearance of lymphocytic choriomeningitis virus (LCMV) in mice (48). Moreover, IL-10 blockade was shown to enhance the magnitude and quality of TH1 responses sufficient to reduce vaccination/boost frequency from three vaccine doses to only one dose (49). Indeed, αIL-10R1 has also been used in combination with BCG to induce anti-cancer immunity which protected mice from bladder cancer (50). Thus, these combined studies highlight the potential for IL-10 modulation as a host directed therapy during vaccination to improve protective immunity against a variety of different disease models, including TB, and we postulate that studies of IL-10 modulation during initial vaccination should be revisited as a mechanism to boost host protective immunity with vaccines that have known diminished long-term protection efficacy, such as BCG. Protective immunity generated by BCG vaccination wanes over time in humans and in most experimental animal models (37, 38, 51–53). Interestingly, BCG vaccination induces strong TH1 responses (54–57), and both *M.tb* infection and BCG immunization induces effector T cells (57–62). However, long lived central memory T cells are associated with superior efficacy of experimental TB vaccines (63–68). In contrast to many short-term microbial infections where a rapid effector T cell response can resolve infection, BCG must stimulate long-term memory immunity to maintain effective control of a recent or a latent *M.tb* infection, often in the absence of sterilizing immunity. Adding additional complexity, BCG also induces IL-10 production which diminishes long-term antigen specific TH1 immune responses (55, 69-72). In this current study, we determined immune responses to αIL-10R1/BCG vaccination at week 1 to observe the immediate impact of IL-101R blockade, and at week 7 post vaccination to characterize the immune status in the lung at the time of *M.tb* challenge. At week 1, BCG vaccination alone increased IL-10 in the lung, which remained high 7 weeks later. Interestingly, αIL-10R1/BCG vaccination reduced IL-10 and led to a concomitant reduction in pro-inflammatory cytokines in the lung. The reduction in pro-inflammatory cytokines was independent of BCG burden as we did not detect BCG in the lungs, and it was accompanied by a modest increase in central and effector memory CD4^+^ and CD8^+^ T cell subsets in the lungs, with enhanced antigen specific TH1 and TH17 responses. Thus, a single αIL-10R1/BCG cocktail dose was capable of modifying the local lung environment immediately following vaccination.

IL-10 is a pleotropic cytokine secreted by various immune cells, with innate cells being a source of IL-10 production during the early phase of BCG vaccination (73). Innate cells can also respond to IL-10 and alter their function (74). Thus, the immediate impact of IL-101R blockade during BCG vaccination may reflect changes in innate cell function including enhanced and prolonged antigen presentation, resulting in altered T cell priming and generation of long-lived T cell responses (27, 75). Indeed, IL-10 from non- B or −T cell sources have been shown to regulate dendritic cell driven TH1/TH2 responses *in-vivo* (75), where, similar to our findings, early IL-10 blockade enhanced TH1/TH17 responses associated with accelerated fungal clearance in mice (76).

Our studies extend observations by Pitt *et al.*, where αIL-10R1 treatment given at the time of BCG vaccination, and for an additional 6 weeks thereafter, promoted antigen specific TH1/TH17 immunity in the lungs, and reduced *M.tb* burden, defining an association between enhanced protection and TH1/TH17 responses (28), However, the extended period of αIL-10R1 treatment (for 6 weeks post vaccination) could not fully separate the timing of the negative influence of IL-10 on the early generation of long-term protective immunity, from effector functions. Our single-dose αIL-10R1/BCG vaccine strategy confirms that IL-10 can negatively impact the initial generation of long lasting protective immunity against *M.tb* infection. Given the short half-life of αIL-10R1/IgG1, a direct pleotropic influence of αIL-10R1 at week 7 post vaccination (our selected time for *M.tb* challenge) was unlikely (77).

The CBA/J mouse strain, the strain background for our studies, has limited protection afforded by BCG (41), making it an ideal WT strain to determine if the αIL-10R1/BCG cocktail can improve protective efficacy and identify potential mechanisms of action. *M.tb* infection of αIL-10R1/BCG vaccinated CBA/J WT mice had a sustained enhanced control of *M.tb* infection and extended lifespan beyond 50 weeks. In contrast, BCG vaccination alone failed to extend survival relative to non-vaccinated CBA/J WT mice. A single dose of αIL-10R1/BCG vaccine was also associated with reduced lung inflammation as determined by histopathology, and reduced pro-inflammatory cytokines in the lung at week 4 and 8 post *M.tb* infection. Pro-inflammatory cytokine production can depend on the bacterial load, yet both BCG and α-IL-10R1/BCG vaccinated mice had similar *M.tb* burden at week 4 post infection, suggesting that α-IL-10R1/BCG vaccination generates an immune response that is less inflammatory, identifying a potential causal relationship between reduced inflammation at the time of vaccination and the generation and prolonged maintenance of IL-17/IFN-γ producing central memory T cells, as described by others ((43). An inflammatory environment can alter the development of effector and/or memory response generating short-lived effector cells (78, 79). Thus, our results support the concept that a strong inflammatory response regulates T cell sensitivity, proliferation and migration of both effector and established memory T cells populations (80, 81).

Our findings in WT mice treated with αIL-10R1/BCG were corroborated by similar studies in IL-10 knockout mice on the same CBA/J mouse strain background (22). BCG immunized IL-10^−/−^ CBA/J mice had superior control of *M.tb* infection and significantly reduced lung immunopathology for up to 50 weeks post infection. Interestingly, both non-immunized and BCG immunized IL-10^−/−^ mice had 100% survival for at least 50 weeks post infection (study end point), albeit with different immunopathology scores suggesting a possible split in survival much later. Modulation of IL-10R1 at late stages of *M.tb* infection is already known to improve outcomes (21). However, the use of a non-conditional knockout model, while complementary, cannot spatially separate this from the influence of IL-10R1 during T cell priming but it can substantiate IL-10 as the primary driver of our observed phenotypes.

Overall, our studies indicate that temporal and spatial blocking of IL-10R1 is sufficient to generate long-term protective immunity against *M.tb* infection in the relatively susceptible CBA/J mouse strain. Our studies identify IL-10, or its downstream effector functions, as a putative target for the development of improved vaccines for TB, and identify a single dose vaccine strategy against *M.tb* that can result in long-term protective immunity and reduced TB disease.

## Acknowledgements

We thank the animal resources personnel and Biosafety Level 3 Program at both The Ohio State University and Texas Biomedical Research Institute. The following reagent was obtained through BEI Resources, NIAID, NIH: *Mycobacterium tuberculosis*, Strain H37Rv, Culture Filtrate Proteins (CFP), NR-14825.

## Funding

This work was supported by The Ohio State University (OSU) College of Medicine, OSU Public Health Preparedness for Infectious Disease (PHPID) pilot grant program (JT), Texas Biomedical Research Institutional funds (JT), and by the Robert J. Kleberg, Jr. and Helen C. Kleberg Foundation (JBT).

## Conflict of Interest

Authors declare non-conflict of interest.

## Notes

### Competing Interest Statement

The authors have declared no competing interest.

